# Molecular Dynamics Simulation of a Model Brain Membrane with and without a 14-3-3 tau Protein

**DOI:** 10.1101/2021.07.31.454604

**Authors:** Jorge A. Arvayo-Zatarain, Fernando Favela-Rosales, Angel D. Reyes Figueroa, Pavel Castro-Villarreal, Claudio Contreras-Aburto

**Affiliations:** Facultad de Ciencias en Física y Matemáticas, Universidad Autónoma de Chiapas, Carretera Emiliano Zapata, Km. 8, Rancho San Francisco, C. P. 29050, Tuxtla Gutiérrez, Chiapas, México; Departamento de Investigación, Instituto Tecnológico Superior Zacatecas Occidente, Ave. Tecnológico 2000, 99102 Sombrerete, Zacatecas, Mexico; Department of Chemistry, Western University, 1151 Richmond St, London ON, CA. N6A 5B7; Facultad de Física, Universidad Veracruzana, Circuito Aguirre Beltrán S/N, 91000 Xalapa, Veracruz, México

## Abstract

An unbalanced composition of lipids and proteins in brain membranes is related to the appearance neurodegenrative diseases and recent investigations show that the 14-3-3 tau protein might relate to some of these diseases. This article reports results from a coarse-grained model brain membrane with and without a 14-3-3 *τ/θ* protein inside the membrane. We investigated the symmetrized partial density, thickness, diffusion coefficients, and deuterium order parameters of the membrane with and without protein. We observe a slight increase in heads and linkers in the symmetrized partial density of the membrane with the protein inserted and higher values of the deuterium order parameters for the brain model membrane with protein. We observe a reduction in the diffusion coefficient of the fluid membrane in the presence of the transmembrane tau protein. Our findings show that the protein can modify the structural and dynamical properties of the membrane. This work will serve as a guide for future investigations on the interactions of tau proteins with brain membrane models and their relation to neurodegenerative diseases.

## Introduction

Cell membranes are biological structures composed of a large variety of lipids and proteins. Somen are integral proteins exposed to the cell interior and exterior parts and are called transmembrane proteins. Cell membranes serve as semi-permeable barriers that behave as a two-dimensional fluid at physiological temperatures that can host many proteins and other types of macromolecules.^1,2^ Lipids in the membranes do not distribute uniformly (i.e., membranes are highly heterogenous media) and their composition varies between different kinds of membranes (cellular or organelles). Inappropriate compositions of lipids and proteins are related to the appearance of many diseases like Parkinson’s, cancer, diabetes, and Alzheimer’s. ^3^ There are proteins linked to the appearance of some of these diseases, e.g., the family of 14-3-3 proteins. The brain is the tissue with the most concentration of these proteins.

The 14-3-3 proteins are phospho-serine/phospho-threonine binding proteins involved in several mechanisms, they interact with kinases and participate in transcriptional processes. There are seven isoforms of this protein commonly labeled with Greek the letters: (*β, γ, ϵ, η, ξ, σ* y *τ*/*θ*). Each isoform consists of nine alpha-helices which form an amphipathic groove that can bind to a protein partner. Because of their incidence in neurological diseases, the interactions of 14-3-3 proteins with brain membranes is a current topic of research^4–6^

One of the essential 14-3-3 binding partners is the isoform tau, a microtubule-associated protein. These proteins have been found in the cerebrospinal fluid of people with various neurological disorders and seem to assemble in some specific disease-lesions of the brain. Researchers found them in the cerebrospinal fluid of patients with dementia of unknown causes, fronto-temporal dementia, and brain tumors. They are also found in NFT’s of Alzheimer’s disease brain sections.^7–18^ and in patients with Parkinson’s disease protein aggregates known as Lewy Bodies.^19–22^

Another important function of the 14-3-3 proteins is the neuroprotective effect in the way of the inhibition of the apoptotic process by multiple mechanisms. This causes cell survival. Also, the 14-3-3 proteins extend their neuroprotective effect by the formation of aggresomes and this facilitates the degradation of misfolded toxic proteins. Recently, has been demonstrated that the accumulation of toxic proteins is one of the main causes of neurodegeneration and the elucidation and study of 14-3-3 proteins and its molecular mechanisms of for example, formation of aggresomes, could be important on the investigation of neuropathologies.^23,24^

Although it is known that 14-3-3 proteins are involved in neurological disorders and the brain is the tissue with most concentration of these proteins, most of the computational studies involve the beta amyloid in comparison of 14-3-3 proteins. ^25–27^ Ingólfsson, et al., ^3^ recently reported a study of a realistic human brain plasma membrane using advances in computational power and molecular dynamics simulation (MD) techniques.

In this work, we simulate using MD a coarse-grained model the interaction of a 14-3-3 protein with/inside a model brain membrane. We report results for a brain plasma membrane constructed on the basis of the composition reported by Ingólfsson et al., adding a 14-3-3 *τ/θ* protein inside the membrane. We investigate the symmetrized partial density, thickness, diffusion coefficients, and deuterium order parameters of the membrane with and without the protein. We observe a slight increase in heads and linkers in the symmetrized partial density of the membrane with the protein inserted and higher values of the deuterium order parameters for the brain model membrane with protein. We observe a reduction in the diffusion coefficient of the fluid membrane in the presence of the transmembrane tau protein. Our findings show that the protein can modify the structural and dynamical properties of the membrane.

## Methods

Using the INSANE program, two membranes were constructed: one with 14-3-3 *τ* isoform inserted and the other without the protein. The neural plasma membrane composition was obtained from Ingólfsson, et al. ^3^ The simulations were performed using the Martini Force Field. ^28,29^ The parameters for lipids, ganglioside and improved cholesterol were described according to Ingólfsson et al, López et al and Melo et al, respectively. The lipid composition is based in the membrane reported by Ingólfsson et al.^3^ but is not at the same proportions as the reported by them. The details of the composition of the model brain membrane are shown in tables 1 and 2, and in the pie charts of figure 1.

**Table 1:** Total Composition of Membrane without 14-3-3 Tau.

| LIPID | TOTAL |
| --- | --- |
| PC | 388 |
| PE | 364 |
| SM | 118 |
| GM1/GM3 | 28 |
| CEREBROSIDES | 77 |
| CERAMIDES | 15 |
| LPC/LPE | 10 |
| DAG | 8 |
| PS | 130 |
| PI | 66 |
| PA | 4 |
| PIPs | 15 |
| CHOL | 1052 |
| W | 21827 |
| NA | 394 |
| Cl | 87 |

**Table 2:**
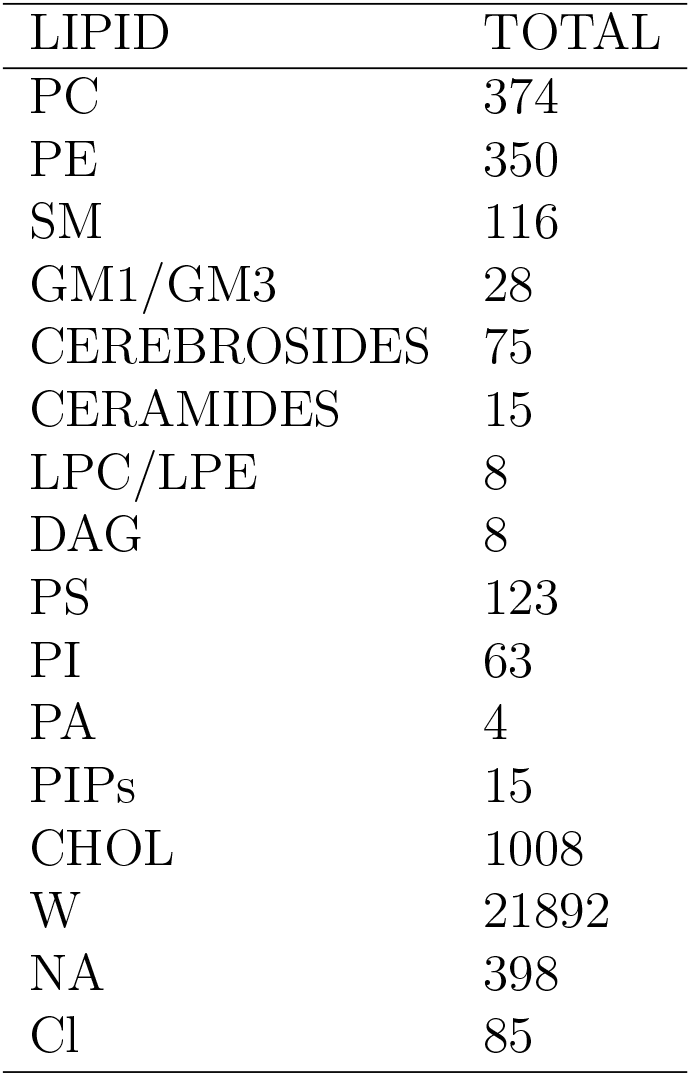
Total Composition of Membrane with 14-3-3 Tau.

| LIPID | TOTAL |
| --- | --- |
| PC | 374 |
| PE | 350 |
| SM | 116 |
| GM1/GM3 | 28 |
| CEREBROSIDES | 75 |
| CERAMIDES | 15 |
| LPC/LPE | 8 |
| DAG | 8 |
| PS | 123 |
| PI | 63 |
| PA | 4 |
| PIPs | 15 |
| CHOL | 1008 |
| W | 21892 |
| NA | 398 |
| Cl | 85 |

**Figure 1:**
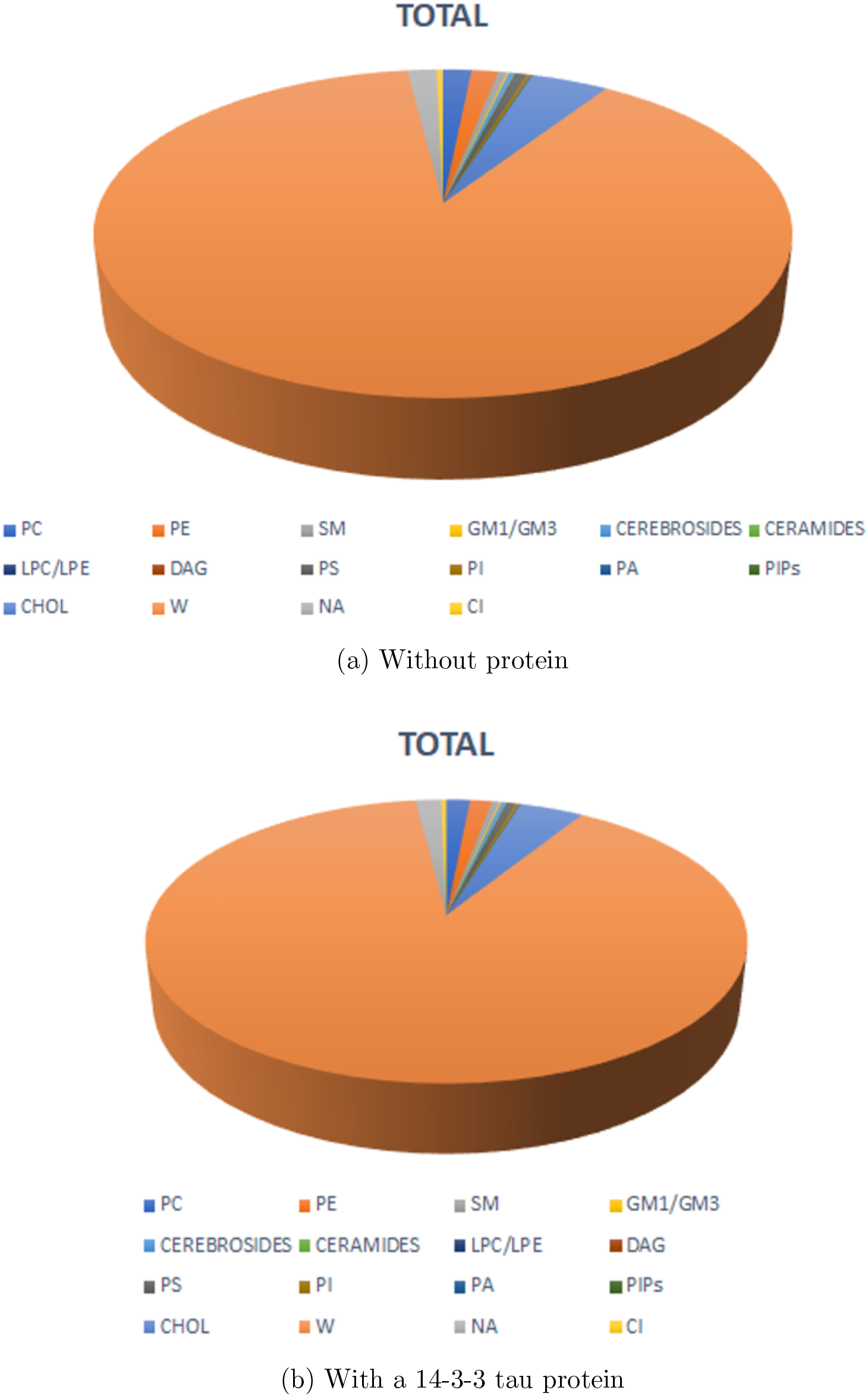
Pie charts representing the lipid composition of the membrane

The simulations were run using GROMACS 2019.4 with a timestep of 10 fs. The simulation box of a model brain membrane without 14-3-3 *τ* contained 2275 lipids, 21892 and 481 mM NaCl. The simulation box of the model brain membrane with 14-3-3 *τ* protein contained 2187 lipids, 21892 waters and 483 mM NaCl. For each bilayer mixture, the composition was adjusted to have a symmetrical composition of each leaflet. The temperature and pressure (310 K and 1 bar of pressure) were controlled with the velocity rescaling thermostat and the Parrinello Rahman barostat. The total time of simulation of the equilibrated system was 9 *µ*s. Figures 2 and 3 illustrate the initial and final configurations on the time interval of 9 *µ*s of the equilibrated systems, with and without the tau protein, respectively.

**Figure 2:**
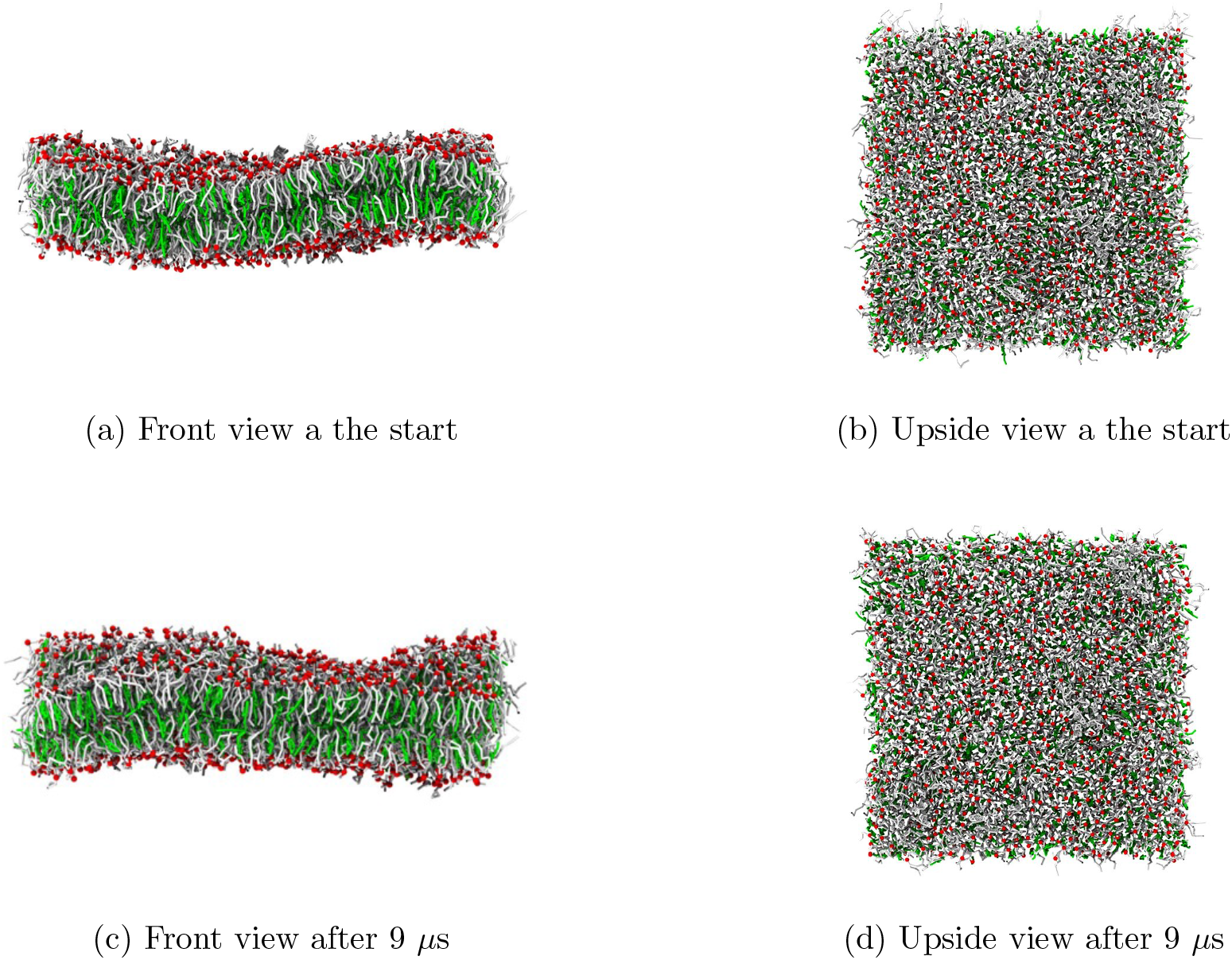
Snapshots of the membrane in equilibrium without protein on a time interval of 9 *µ*s.

**Figure 3:**
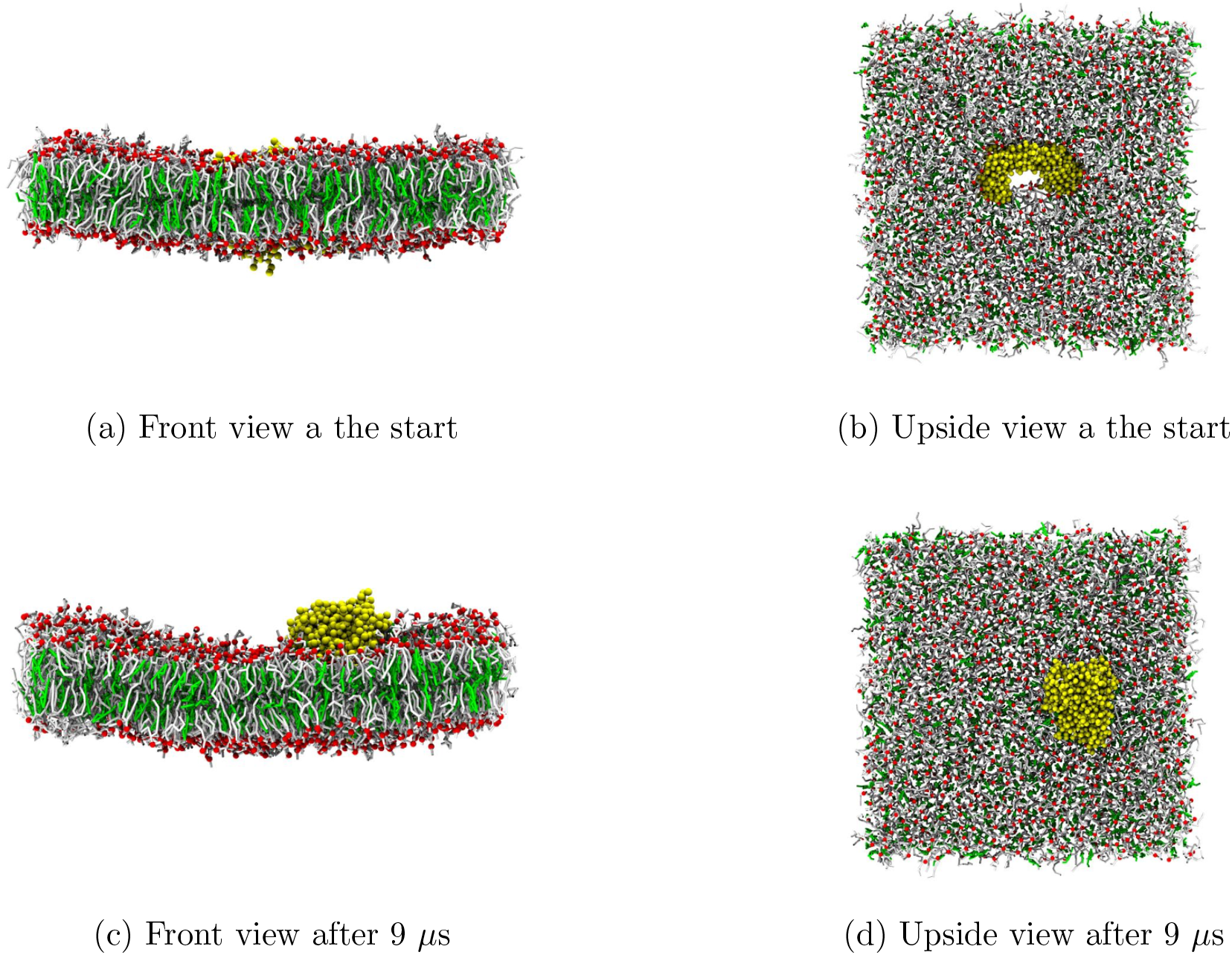
Snapshots of the membrane in equilibrium with a 14-3-3 Tau protein, on a time interval of 9 *µ*s.

The diffusion coefficient was obtained from Einstein relation

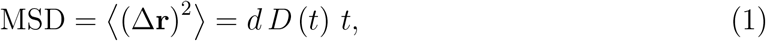

where MSD stands for mean squared displacement, Δ**r** is the displacement vector of a particle occurring on the time interval *t, d* is the dimensionality (*d* = 3 for diffusion in 3D euclidean space, *d* = 2 for diffusion in 2D euclidean space) and *D*(*t*) is the so-called self-diffusion coefficient. The variable *t* is also called the correlation time. The usual diffusion coefficient is the long-time limit of *D*(*t*).

## Results and discussion

The model brain membranes with and without 14-3-3 *τ* were correctly equilibrated at 310 K and 1 bar. From the simulated trajectories we performed the analysis of following properties of the membrane with and without 14-3-3 tau:

1. Density profiles of Heads, Tails and Linkers.
2. Deuterium order parameters of each species.
3. Local membrane thickness
4. Self diffusion of PO4 and ROH.

The results displayed in figure 4 show that there are only slight differences between both density profiles of the tails, heads and linkers. The tail density with 14-3-3 *τ* inserted peaks at about 1200 kg m^*−*3^, heads at (about 400 kg m-3) and linkers, at 350 kg m^*−*3^ and in the model brain membrane the density was 300 kg m^*−*3^. These values are consistent with the results of Ingólfsson et al.

The details of the order parameters of SN1 and SN2 tails are given in tables 3 and 4 for the membrane with and without protein, respectively. The average order parameter of SN1 and SN2 tails are respectively 0.29190725 and 0.3611618133 without protein, while with the protein they are 0.3262190325 and 0.3921093714. We observe an increase of about ten percent in the order parameter due to the presence of the protein and this implies more order of the carbon tails. This explains why the model brain membrane without protein presents more undulations in comparison with the membrane containing the protein inserted, as shown in the snapshots of figures 2 and 3. One cay say that the protein makes the membrane slightly more stiff.

**Table 3:**
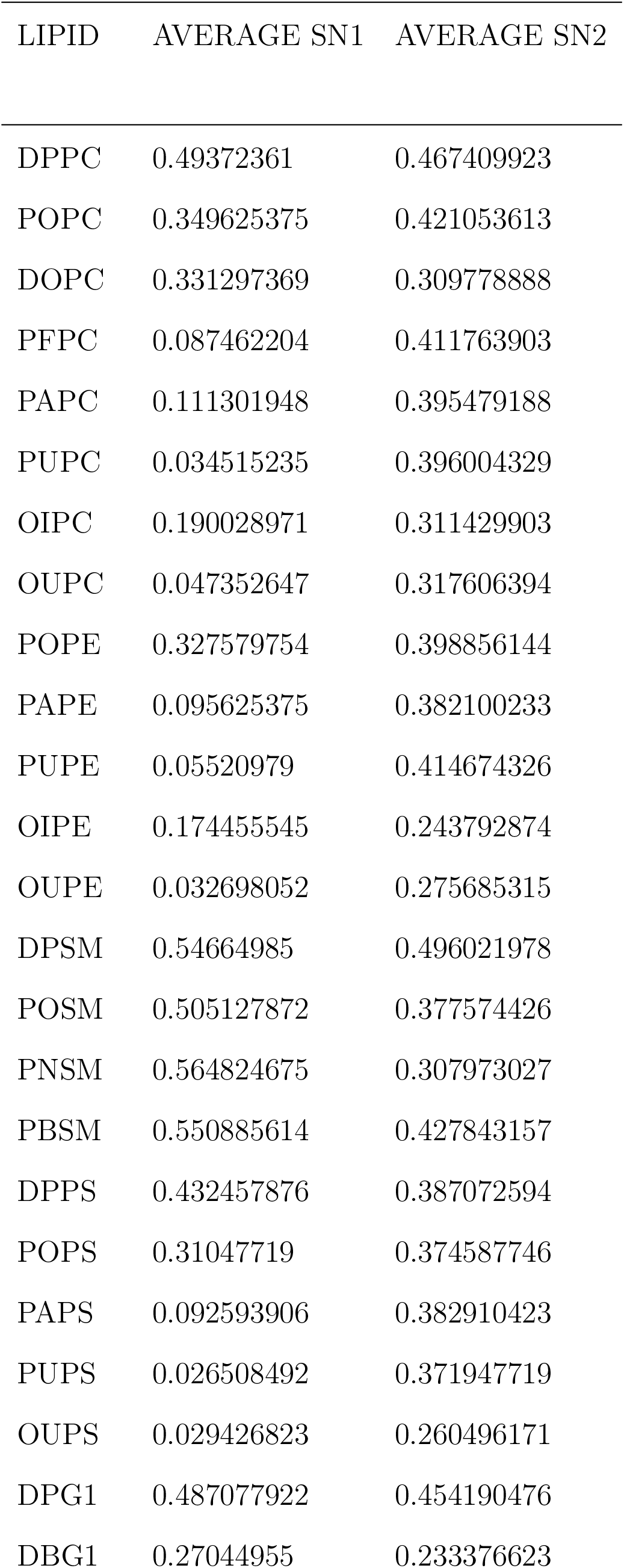
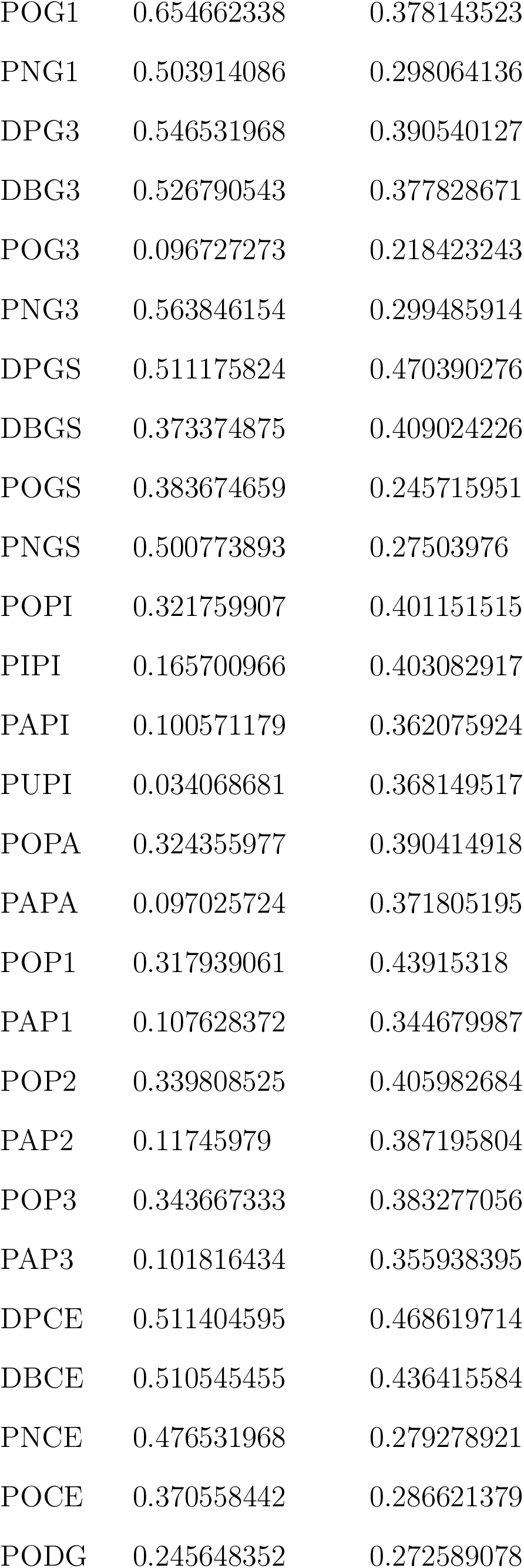
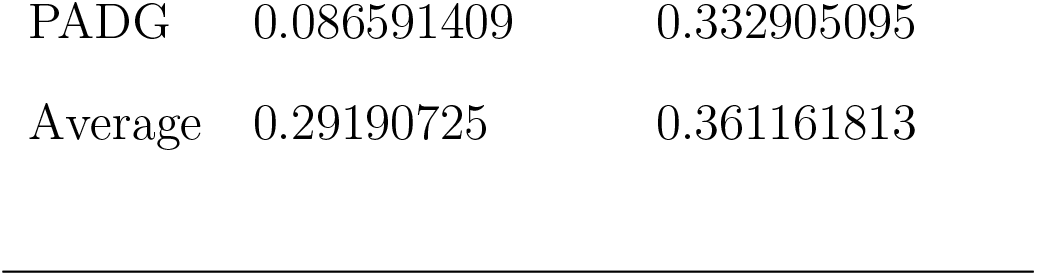
Deuterium order parameters of tails (SN1 and SN2) without Tau.

**Table 4:** Deuterium order parameters of tails (SN1 and SN2) with Tau.

| LIPID | AVERAGE SN1 | AVERAGE SN2 |
| --- | --- | --- |
| DPPC | 0.521338661 | 0.495453547 |
| POPC | 0.37435298 | 0.451484516 |
| DOPC | 0.35506327 | 0.336900433 |
| PFPC | 0.110277389 | 0.452552115 |
| PAPC | 0.111145854 | 0.433031302 |
| PUPC | 0.043435564 | 0.43720979 |
| OIPC | 0.219738262 | 0.331482518 |
| OUPC | 0.053198052 | 0.314061272 |
| POPE | 0.365443889 | 0.429027639 |
| PAPE | 0.105071262 | 0.412958042 |
| PUPE | 0.048997253 | 0.420397935 |
| OIPE | 0.205520147 | 0.295971695 |
| OUPE | 0.049857393 | 0.307829837 |
| DPSM | 0.601566434 | 0.540772228 |
| POSM | 0.531513986 | 0.396740926 |
| PNSM | 0.595387612 | 0.343360639 |
| PBSM | 0.59114985 | 0.464707792 |
| DPPS | 0.474235764 | 0.446102897 |
| POPS | 0.362480853 | 0.432795538 |
| PAPS | 0.102448801 | 0.411544789 |
| PUPS | 0.034364136 | 0.421157842 |
| OUPS | 0.034838412 | 0.305901099 |
| DPG1 | 0.456264735 | 0.475478189 |
| DBG1 | 0.284046287 | 0.174887113 |
| POG1 | 0.614955045 | 0.360814519 |
| PNG1 | 0.511564436 | 0.327770629 |
| DPG3 | 0.492808192 | 0.467549118 |
| DBG3 | 0.552454546 | 0.462795205 |
| POG3 | 0.687015984 | 0.318240759 |
| PNG3 | 0.597289211 | 0.303240559 |
| DPGS | 0.501782884 | 0.484575758 |
| DBGS | 0.479393856 | 0.48571978 |
| POGS | 0.539064269 | 0.391885448 |
| PNGS | 0.406715951 | 0.262788811 |
| POPI | 0.31160706 | 0.393394605 |
| PIPI | 0.195031635 | 0.435511489 |
| PAPI | 0.109324176 | 0.404929737 |
| PUPI | 0.038907093 | 0.370140526 |
| POPA | 0.42853713 | 0.498704296 |
| PAPA | 0.106294955 | 0.398943723 |
| POP1 | 0.372774226 | 0.422955378 |
| PAP1 | 0.105776723 | 0.377894106 |
| POP2 | 0.37044289 | 0.443132534 |
| PAP2 | 0.129152348 | 0.414649684 |
| POP3 | 0.391010989 | 0.472774226 |
| PAP3 | 0.13682018 | 0.436742591 |
| DPCE | 0.545969031 | 0.516066933 |
| DBCE | 0.538099567 | 0.466048701 |
| PNCE | 0.582164336 | 0.30054006 |
| POCE | 0.441255744 | 0.291197136 |
| PODG | 0.313405594 | 0.363811522 |
| PADG | 0.049972361 | 0.068100233 |
| Average | 0.326219033 | 0.392109371 |

**Figure 4:**
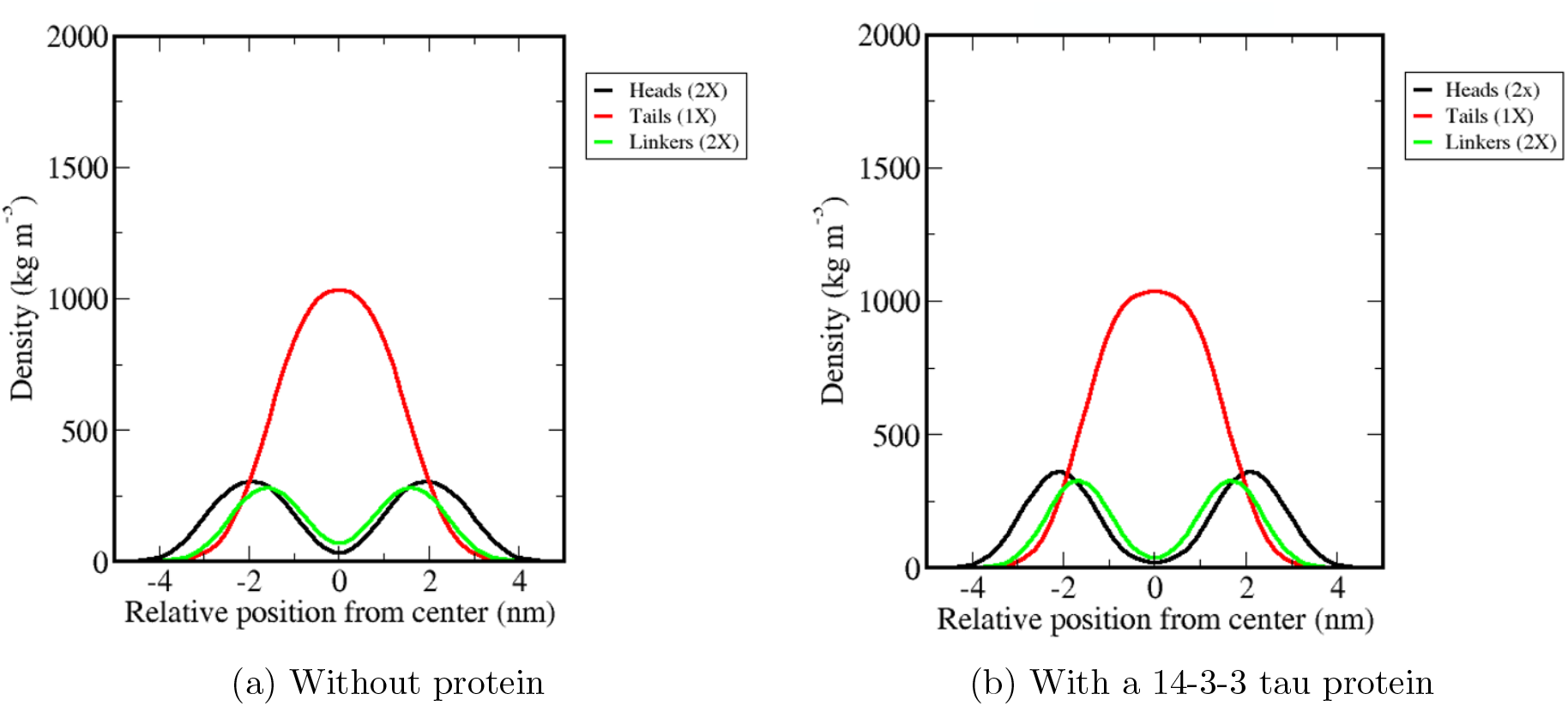
Density profiles along the *z*-direction

Figure 5 shows the dynamics of the membrane thickness averaged on 100 nanoseconds of each of the microseconds of the simulation run, with and without protein. We observe that the thickness exhibits smaller fluctuations when the protein is present. In a sense, the protein stabilizes the model brain membrane.

**Figure 5:**
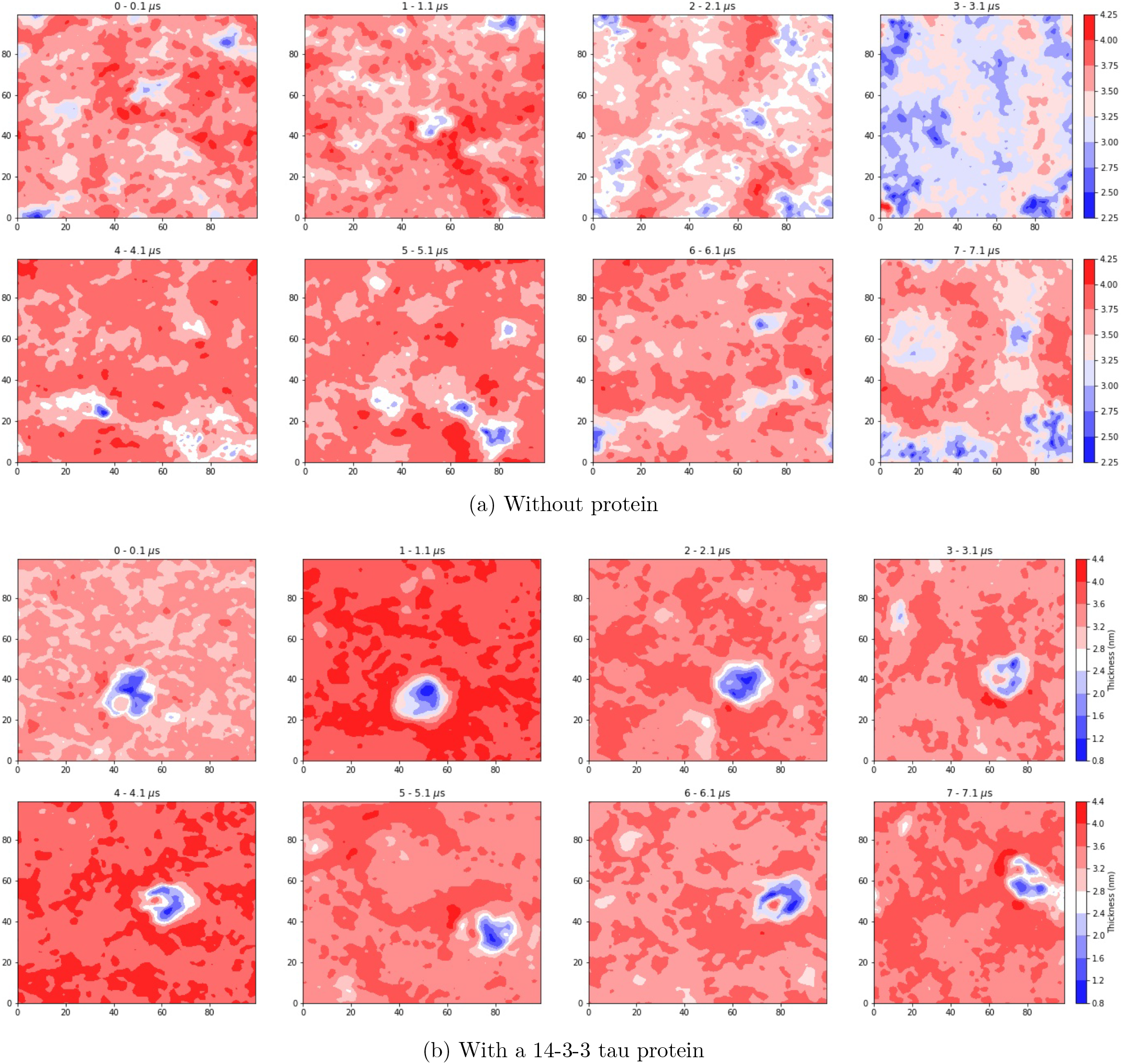
Temporal evolution of membrane thickness

Figure 6 displays de average MSD self-diffusion coefficient *D*(*t*) of PO4 and ROH in 3D and 2D (XY plane) spaces. The average is taken in different regions of space as indicated in the figure. The graphs labeled PO4 and ROH are the self-diffusion coefficients averaged over all the corresponding group of molecules of PO4 and ROH molecules in the simulation box, respectively, without the protein inserted. The graphs labeled PO4*<*5nm and ROH*<*nm are the self-diffusion coefficients averaged over all the corresponding group of molecules located inside a sphere of radius 5nm around the center of the protein. The graphs labeled PO4*>*5nm and ROH*>*nm are the self-diffusion coefficients averaged over the groups of molecules located at the exterior of a sphere of radius 5nm around the center of the protein in the simulation box. The PO4 and PO4*>*5nm graphs are practically indistinguishable form each other, and the same can be said about the ROH and ROH*>*5 graphs. This means that at the level of the dynamics, the influence of the protein is limited to a cutoff radius of about 5nm. On analyzing the numerical values of the self diffusion coefficient in the long time regime one observes that the effect of the protein is to reduce the diffusion coefficient in an amount of 8 to 10 per cent with respect to the value in absence of the protein. Furthermore, note that the obtained value of the diffusion coefficient is of the order of 25 *µ*m^2^ · ns^*−*1^. The authors of the Martini force field mention that “to compare the diffusion coefficient obtained from a Martini simulation to a measured one, a conversion factor of about 4 has to be applied to account for the faster diffusion at the CG level due to the smoothened free energy landscape”. Taking into account this scaling factor, our results for the diffusion coefficient are in good agreement with typical values of lipid diffusion coefficient obtained experimentally.

**Figure 6:**
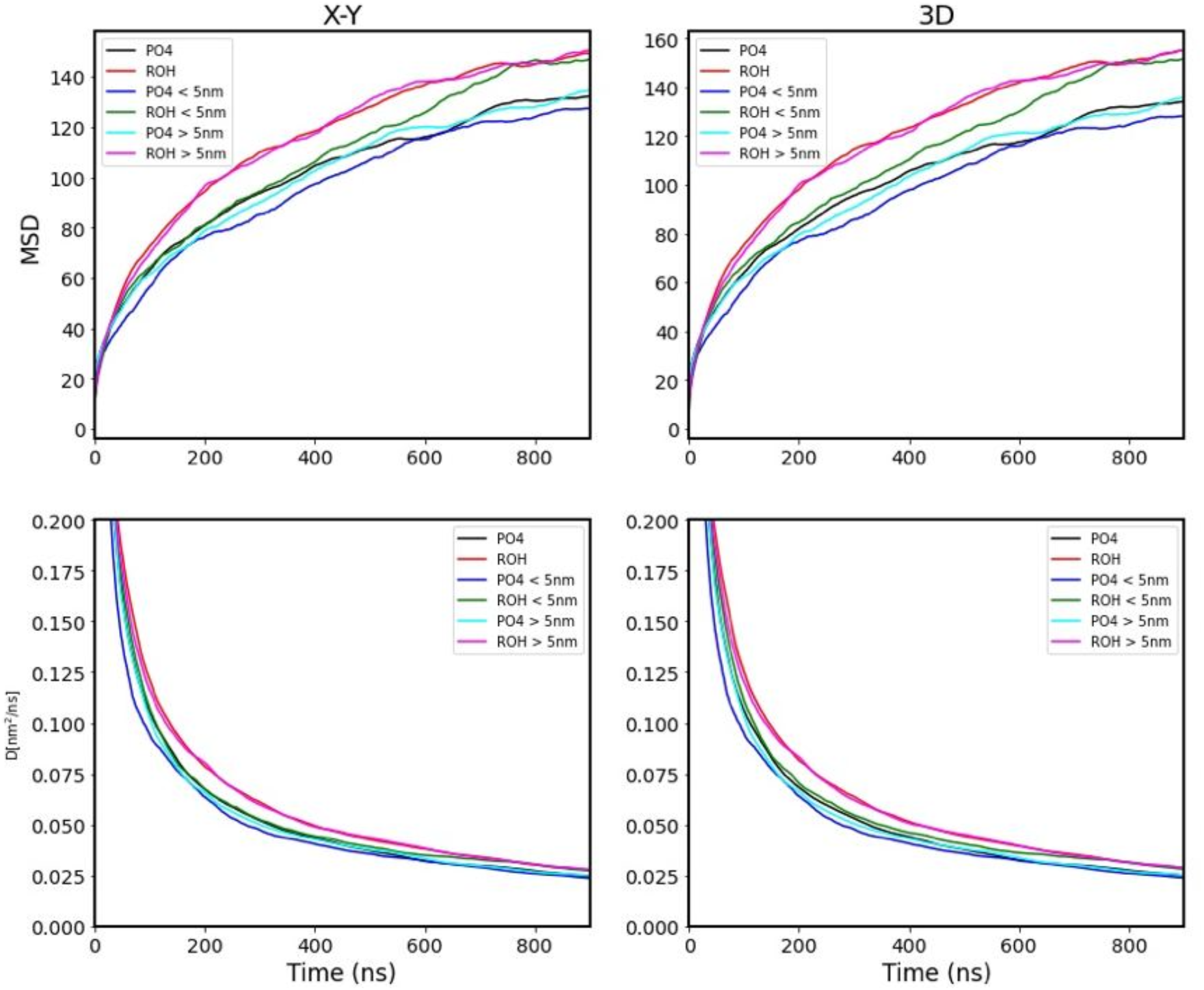
Above: Mean squared displacement along the XY plane and in the 3D space. Below: Self-diffusion coefficient on the XY plane and in the 3D space.

## Conclusions

In this work, we correctly built two realistically model brain membranes: one with 14-3-3 protein inserted and the other without 14-3-3 *τ* inserted, with the primary objective to observe the influence of the protein inside the model brain membrane.

We obtained that the protein reduces the diffusion of the closer lipids, reduces thickness fluctuations, and increases the order of the SN1 and SN2 tails. In the analysis of the partial density of the membrane, we observe in the model brain membrane with protein inserted a slight increase in the peaks of lipid heads and linkers, compared to the model brain membrane. Also, in the order parameters, we can observe higher values in the model brain membrane with the protein inserted than the membrane without protein. We can conclude that the presence of 14-3-3 tau slightly modifies the undulations of the membrane and makes the membrane more stable to thickness fluctuations.

There are many interesting questions about the role of the 14-3-3 protein in model brain membranes that we should tackle in the near future. For example, investigate the influence of a finite protein concentration on the properties of the membrane.

## Acknowledgement

J.A.A.-Z. thanks CONACYT for financial support through the program Estancias Posdoctorales Nacionales, and Facultad de Ciencias en Física y Matemáticas UNACH. We thank ACARUS-Unison and LARCAD-UNACH for access to computing facilities.

